# The repertoire of testicular extracellular vesicles cargoes and their involvement in inter-compartmental communication required for spermatogenesis

**DOI:** 10.1101/2021.01.08.426002

**Authors:** Kathleen Hoi Kei Choy, Sze Yan Chan, William Lam, Jing Jin, Tingting Zheng, Sidney Siubun Yu, Weiping Wang, Linxian Li, Gangcai Xie, Howard Chi Ho Yim, Hao Chen, Ellis Kin Lam Fok

**Author notes:** These authors contributed equally to the work (Kathleen Hoi Kei Choy, Sze Yan Chan, William Lam). To whom correspondence should be addressed to Ellis Kin Lam Fok and Chen Hao.

## Abstract

Extracellular vesicles (EVs) secreted by the epididymis and prostate are involved in sperm functions and epigenetic inheritance. However, the EVs in the testis remains unexplored. Here, we have established a testis dissociation protocol that allows the isolation of testicular EVs by minimizing the disruption of fragile sperm cells. We showed that testicular EVs were specifically and efficiently uptaken by somatic cells and germ cells in both interstitial space and seminiferous tubules compartments, including the spermatozoa. We profiled the proteome of testicular EVs and probed the cell types that release them. Moreover, we sequenced the small RNAs cargos of testicular EVs and identified sets of small non-coding RNAs that were overlooked in the testis transcriptome. Selected miRNA candidates in testicular EVs were found in sperm RNA payload and demonstrated specific resistance towards ribonuclease A independent of the vesicle membrane. Small molecule inhibition of EVs secretion increased the apoptosis of germ cells via inter-compartmental communication. Together, our study has provided valuable resources on the cargoes of testicular EVs and revealed the inter-compartmental communication that regulates spermatogenesis and may implicate in paternal inheritance.

## INTRODUCTION

Spermatogenesis requires precisely orchestrated cellular communication networks. This involves mechanisms that are ubiquitously used and shared in other organ systems as well as the mechanisms that are unique to the testis ^1^. Autocrine and paracrine signalling is a common cell-cell communication in various systems. The testicular microenvironment contains cytokines, peptides, metabolites, hormones and growth factors involved in stem cell homeostasis, cell differentiation and cell survival during spermatogenesis ^2–9^.

Cell junction is another means involved in cell-cell communication. The blood-testis barrier formed by the tight junctions of adjacent Sertoli cells is a unique tight junction required for the compartmentalization of the seminiferous tubules. Other testis-specific cell junctions formed at the Sertoli-Sertoli interface and Sertoli-germ cell interface, also known as the basal ectoplasmic specialization (ES) and apical ES respectively, are essential for various stages of germ cell development, the phagocytosis of cytoplasmic bodies during spermiogenesis and the release of sperm to the lumen after the spermiogenesis ^8,10,11^.

Another unique cell-cell communication in the testis is the intracytoplasmic bridges formed in sister spermatogonia after rounds of mitosis. A recent study has demonstrated that the intracytoplasmic bridges can exchange macromolecules such as proteins and RNAs. Besides, the fragmentation of intracytoplasmic bridges may be involved in cell fate determination of the spermatogonial stem cell population ^12^.

EV is an important mechanism of cell-cell communication. The EVs can be classified into three classes according to their size. The largest size of EVs, typically >1000 nm, is known as the apoptotic bodies. These apoptotic bodies are usually found by blebbing of the apoptotic cells. Microvesicles are EVs with sizes ranging from 100-1000 nm. They are formed by budding and shedding of the plasma membrane. The smallest size of EVs is exosome, with sizes ranging from 50-100 nm. They are produced inside the cells and released via exocytosis ^13^. These EVs carry various cargos, including genomic DNA, mRNA, miRNA and proteins that can be uptake by target cells. Accumulating evidence suggests that proteins on the EVs may be associated with cell recognition and thus target a specific cell type. Furthermore, the delivery of cargos could regulate the cellular processes of the target cells ^14–16^, indicating that the EVs are an important and effective way of cell-cell communication.

In the male reproductive tract, EVs from the epididymis and prostate have been characterized. The epididymosomes consisted of two major classes: the CD9+ epididymosomes and ELSPBP1-enriched epididymosomes. The CD9+ epididymosomes transfer their protein cargos to the spermatozoa and regulate sperm maturation. ELSPBP1-enriched epididymosomes bind preferentially to the dead spermatozoa and quench the reactive oxygen species that may otherwise exert adverse effects on sperm maturation ^17^. Besides, epididymosomes also convey RNA to spermatozoa. By comparing the small RNA transcriptome of spermatozoa before and after epididymal maturation, it was shown that the number of miRNAs remains consistent while a marked decrease in piRNA and a significant increase in tRNA fragments (tRFs) is observed after maturation ^18–20^.

Similar to the epididymosomes, the protein cargos in prostasomes are also delivered to the spermatozoa and implicate in the survival and motility of sperm via calcium-dependent signalling ^21^. Although the prostasome contains DNA, coding and regulatory RNAs with potential modulatory functions, evidence on the transfer of these nucleic acids to the spermatozoa remains scarce.

Surprisingly, although the testis is essential for sperm production, the presence of EVs in the testis and their involvement in spermatogenesis remain largely unknown. Of note, since spermatozoa is transcriptionally inactive, recent studies have postulated that the testicular microenvironment may convey RNA to spermatozoa through EVs ^22^. However, the testicular EV is poorly characterized. Here, we have developed a one-step testis dissociation method for the preparation of single testicular cell suspension. We have isolated and characterized the testicular EVs and showed that the testicular EVs were uptake by somatic cells and germ cells, including the sperm. We further profiled the protein and small RNA cargos of testicular EVs and probed for the cells that release the testicular EVs. Intriguingly, some selected miRNA candidates demonstrated exceptional stability against RNase treatment and were found in the sperm RNA payload. Our study has provided new insights into the intercellular communication in the testis that has broad implications in spermatogenesis and paternal epigenetic inheritance.

## RESULTS

### Establishment of one-step testis dissociation method

The isolation of EVs requires the dissociation of tissue into a single cell suspension with minimal cell disruption in order to minimize the contamination from membranous organelles. To achieve these, we compared the widely adopted two-step double enzyme digestion protocol, which utilized collagenase and trypsin ^23,24^, with a one-step protocol that utilized a non-mammalian, non-bacterial cell dissociation buffer (Fig. S1A). Single testicular cell suspension can be obtained in both methods (Fig S1), but the one-step method demonstrated a significantly higher cell yield and comparable cell viability with less hands-on time (Fig. S1B-E). The one-step testis dissociation protocol also allows the isolation of spermatogonia by magnetic activated cell sorting against Thy1 with a significantly higher yield (Fig. S1D), suggesting the cell surface antigen was better preserved. To further estimate the cell composition, we analyzed the DNA ploidy in the testicular single-cell suspension by flow cytometry. The diploid:tetraploid:haploid ratio in the two-step protocol was 1:2:2 compared to 1:1:8 in the one-step protocol (Fig. S1F-G). This result suggests that the one-step method preserves the haploid sperm cell population which was lysed in the two-step method and maybe a source of organelle contamination in testicular EVs isolation.

### Testicular EVs contains microvesicles and exosomes

We then isolated the extracellular vesicles from the supernatant of the testicular single-cell suspension by differential centrifugation and commercially available membrane affinity column, two widely-adopted methods for EV isolation ^25^. Testicular EVs isolated from double-enzymes digestion revealed a plethora of debris surrounding the EVs (Fig. S2A), suggesting potential contamination of organelles. Consistent with previous reports, differential centrifugation of supernatant from one-step testis dissociation at 10,000 g and 100,000 g isolated microvesicles and exosome that showed the hallmark cup-shape morphology of EVs, with an average size of 700 nm and 300 nm respectively (Fig. S2B-C). We have also attempted to isolate EVs from conditioned medium obtained from mouse primary spermatogonial culture, spermatogonial cell lines C18-4 and GC1-spg, and the Sertoli cell line TM4 (Fig. S2D). Intriguingly, unlike the testicular microenvironment, the cultured cells released trace numbers of EVs with variable sizes.

In corroboration with the differential centrifugation preparation, testicular EVs isolated by membrane affinity column also revealed two distinct peaks at 600 nm and 100 nm with similar cup-shaped morphology (Fig. 1A-B). Testicular EVs prepared by both methods expressed EV markers CD81 and CD63 and showed a negligible level of Golgi markers (Fig. 1C). Notably, while the testicular EVs obtained by membrane affinity column showed a low level of calnexin, an endoplasmic reticulum marker used as a negative control in EV preparation ^26^, its level was markedly higher in testicular EVs prepared by differential centrifugation (Fig 1C). These results indicate the presence of exosomes and microvesicles in testicular EVs. Since relatively pure testicular EVs with minimal organelle contamination can be obtained by membrane affinity column isolation after one-step testis dissociation, we have adopted this protocol in subsequent characterization experiments.

**Fig. 1.**
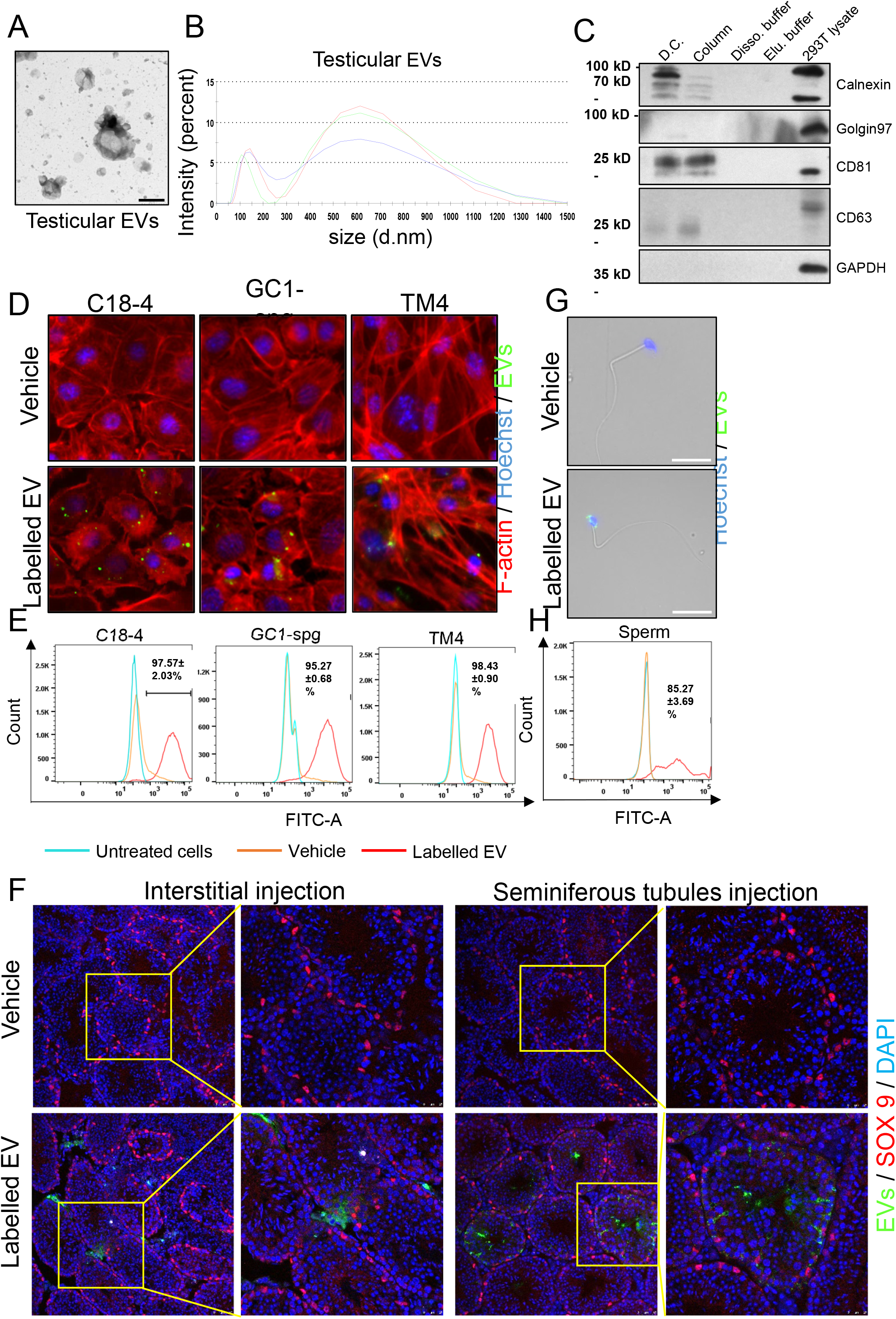
Isolation and characterization of testicular extracellular vesicles. (A) Representative transmission electron microscopy images of testicular microvesicles and exosomes isolated by commercially available affinity columns. Scale bar: 500 nm. (B) Size distribution of testicular EVs isolated from one-step testis dissociation followed by affinity column as determined by dynamic light scattering. (C) Representative Western blot analysis of EV markers CD81, CD63, Golgi marker Golgin 97 and endoplasmic reticulum marker Calnexin in testicular EVs isolated by indicated methods. (D) Representative immunofluorescent images of spermatogonia cell lines (C18-4 and GC1-spg) and Sertoli cell line (TM4) incubated with PKH67 (FITC) labelled testicular EVs for 24 h. The cells were stained for F-actin (phalloidin, red) and nucleus (Hoechst 33342, blue). (E) Representative histograms of testicular EVs uptake in C18-4, GC1-spg and TM4 cells analyzed by flow cytometry. (F) Representative fluorescent images of the uptake of PKH67 labelled testicular EVs in interstitial space (Left panel) and in seminiferous tubules (Right panel). Scale bar: 100 μm. (G) Representative images of sperm incubated with PKH67 (FITC) labeled testicular EVs for 24h. Nucleus were stained by Hoechst 33342. Scale bar: 20 μm. (H) Representative histograms of testicular EVs uptake in sperm analyzed by flow cytometry.

### Uptake of testicular EVs by Sertoli cells and germ cells

EV is an effective way for cell-cell communication. We next sought to investigate the type of cells in the testis that uptake the testicular EVs. We labelled the testicular EVs with PKH fluorescence dye and examined the uptake of fluorescence signals in cell lines and in mouse testis. Our results showed that treating C18-4, GC1-spg and TM4 cell lines with labelled testicular EVs results in the punctated fluorescence signals inside the cells (Fig. 1D). The uptake of labelled testicular EVs in testicular cell lines increased in a time and dose-dependent manner (Fig. S3). Under optimal uptake conditions, flow cytometry analysis confirmed more than 95% of cells in each cell line were fluorescent positive (Fig. 1E), indicating that the tested cell lines were highly efficient in taking up the testicular EVs. Interestingly, compared with the testicular EVs, these cell lines uptake markedly fewer EVs isolated from the conditioned medium of 293 cell line (Fig. S4), suggesting that efficient uptake was specific to the testicular EVs.

In mouse testis, injection of labelled testicular EVs into the interstitial space resulted in marked uptake by interstitial cells and Leydig cells (Fig. 1F). Intriguingly, the fluorescence signals were also observed in the seminiferous tubules, suggesting that the testicular EVs were permeable to the basement membrane and the blood-testis barrier and may be involved in inter-compartment communication. Similarly, when the labelled testicular EVs were injected into the seminiferous tubules, a marked uptake was observed in both the Sertoli cells and various stages of germ cells, including the spermatogonia located at the basal compartment (Fig. 1F). These results support the notion that the testicular EVs are permeable to the blood-testis barrier, although the possibility that the labelled EVs were uptaken and trafficked inside the Sertoli cells and then delivered to the spermatogonia cannot be excluded.

To further investigate the delivery of testicular EVs to spermatozoa, we treated isolated sperm with labelled testicular EVs. We observed the uptake of testicular EVs by more than 84% of sperm and the signals were mainly observed in the head region (Fig. 1G-H). Together, our results suggest the involvement of testicular EVs in the communication between the Sertoli cells and germ cells, including the spermatozoa.

### Proteome of testicular EVs

The protein of EVs may regulate the delivery and modulate the cellular processes in the targeted recipient cells. To profile the proteins carried inside the EVs or on the EV membrane, we examined the testicular EVs proteome by liquid chromatography-mass spectrometry (LS-MS/MS). We have identified 553 proteins in two independent EV preparations (Supplemental Dataset S1). Consistent with our previous finding (Fig. 1C), despite the low abundance, EV markers CD63, CD81 and CD9 were identified in the proteomic study. We next mapped the identified proteins to the corresponding genes and carried out a gene ontology (GO) analysis. The results showed that the vesicle, extracellular membrane-bound organelle and exosome were identified in the GO cellular component analysis (Fig. 2A), suggesting that the testicular EVs shared some degree of similarity in the protein signatures with other systems. Around 50% of the mapped proteins in the testicular EVs were known to be involved in the transport process and 24% possessed substrate-specific transmembrane transporter activity (Fig. 2B-C).

**Fig. 2.**
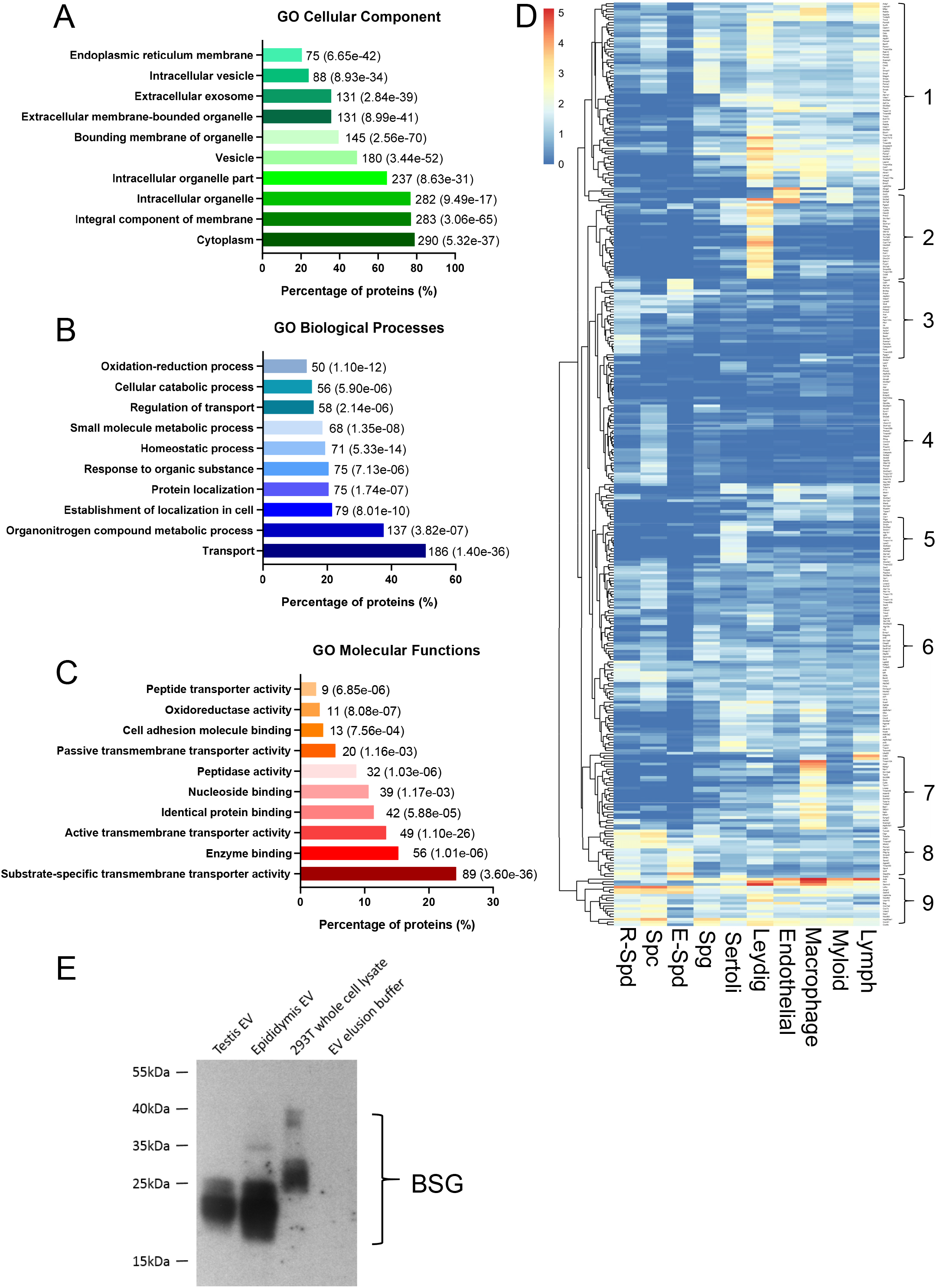
Protein signature of testicular extracellular vesicles. (A-C) Gene Ontology analysis on cellular component (A), biological process (B) and molecular function (C) of proteins identified on/in testicular EVs. The number next to the bar indicates the number of genes that fall in the category with corresponding p value provided in the blanket. (D) Heat map of genes encoding the proteins identified in the testicular EVs and their corresponding expression in indicated types of testicular cells according to single-cell sequencing analysis. Cluster 1 - somatic cell enriched; Cluster 2 - Leydig cell enriched; Cluster 3 - post meiotic germ cell enriched; Cluster 4 spermatocyte enriched; Cluster 5 - Sertoli cell enriched; Cluster 6 - Spermatogonia enriched; Cluster 7 - macrophage enriched; Cluster 8 - germ cell enriched; Cluster 9 commonly expressed. (E) Western blot analysis of candidate protein Basigin (BSG) identified in testicular EVs. EV isolated from the epididymis was used as a control.

We have also sought to probe for the testicular cell types that release the testicular EVs. To do this, we matched the list of identified proteins with the corresponding gene expression from the single-cell sequencing dataset ^27^. We identified 9 clusters of proteins commonly or specifically expressed in various stages of germ cells and various types of somatic cells found in the testicular microenvironment (Fig. 2D), suggesting the secretion of testicular EVs by these cell types. Interestingly, while the cells located in the interstitial space modestly uptake the testicular EVs (Fig. 1F), a set of proteins found in the testicular EVs were enriched in the Leydig cells and macrophages (Fig. 2D). An analysis of the mouse phenotype database revealed that among the top 50 proteins with the highest number of unique mapped peptides in testicular EVs, 11 of them showed a phenotype related to male subfertility or infertility (Fig. S5), suggesting that the importance of testicular EVs that carry these proteins in spermatogenesis. We have validated the proteomic results by Western blot of a selected candidate protein Basigin (BSG), a membrane protein/receptor involved in germ cell migration and survival that was also found in the epididymosome ^2,17,28,29^ (Fig. 2E).

### RNA profiles of testicular EVs

Next, we explored the RNA species carried by the testicular EVs. RNA was isolated by columns for small RNA after proteinase K and RNase A treatment. Bioanalyzer tracing revealed distinct peaks representing small RNA between 25 and ~200 nucleotides length (Fig. S6A). Since the small RNA such as miRNA and tRFs have been shown to mediate cell-cell communication and paternal epigenetic inheritance ^22,30^, we profiled the small RNA in testicular EVs by RNA-sequencing (Supplemental Dataset S2). Testicular EVs mainly carry small RNA that were mapped to genomic loci that were not annotated for known RNA species (73%). These include reads in repeats, exon, intron and intergenic regions. The seven RNA subtypes, miRNA, piRNA, rRNA, tRNA, sncRNA, snoRNA and Rfam sncRNA contributed to the remaining 24% of the total mapped read (Fig. 3A). The piRNA (17%) and miRNA (4%) represent the dominant RNA subtypes among them. Importantly, we have identified sets of novel piRNA and miRNA in the testicular EV samples. These piRNAs and miRNAs may have been overlooked in previous testis transcriptome studies ^31,32^, likely because of the low abundance of testicular EV RNA in the total testis RNA.

**Fig. 3.**
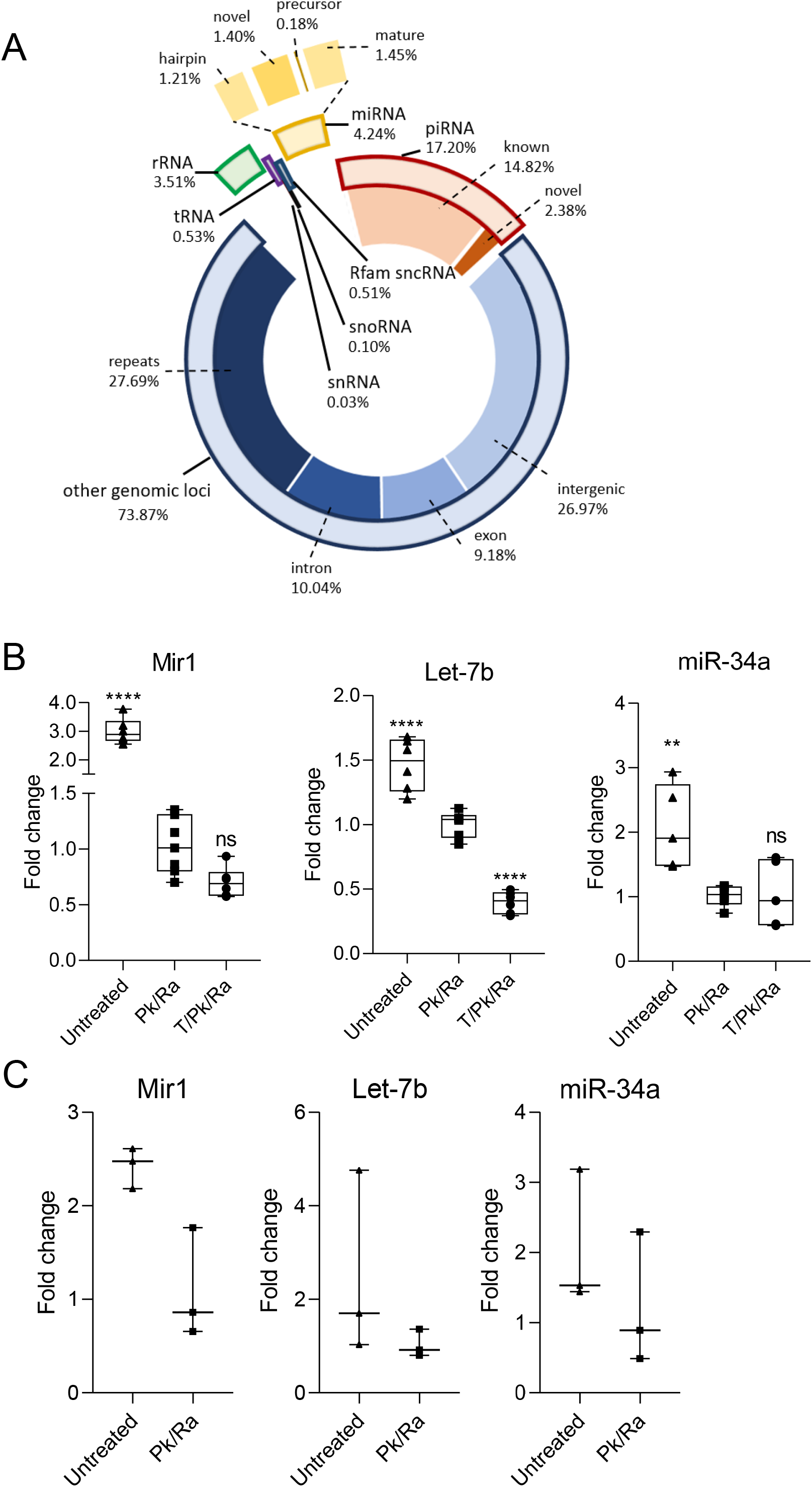
Testicular extracellular vesicles carry miRNA cargoes with differential resistance to RNase in native structure. (A) Diagram showing the distribution of different small RNA species identified in testicular EVs by RNA sequencing and their corresponding percentage counts. Solid bar represent the subclass. (B) Realtime PCR results showing the level of novel Mir1, Let-7b (mature miRNA) and miR-34a (hairpin) in testicular EV samples (B) or sperm (C) without treatment (untreated), or with proteinase K and RNase A treatment (Pk/Ra) or with Pk/Ra treatment in the presence of 1% TritonX-100 (T). Data is presented as mean ± S.D. ** p <0.01, **** p<0.0001, ns - not significant.

Unexpectedly, apart from the rRNA and sncRNA that were mapped to the mouse genome, around 10% of rRNA (0.37% of all tags) and 60% of sncRNA (0.3% of all tags) were classified as bacterial rRNA (both large and small subunits) and transfer-messenger RNA (tmRNA) (Fig. S6B-C), a bacterial RNA molecule with dual tRNA-like and messenger RNA-like properties. This result is consistent with the recent identification of testicular bacteriome in human testis ^33^ and the discovery that rRNA and tRNA are released as the dominant RNA species in the bacterial membrane vesicles ^34^. Together, these findings strongly suggest the presence of microbial-released vesicles in the testicular EVs.

### RNase resistance of testicular EVs miRNA cargos

We then sought to validate the most abundant novel miRNA, Mir1, together with two well-studied miRNAs, miR34a (hairpin) and let-7b (mature). All three miRNAs were identified in testicular EVs samples treated with proteinase K and RNase A, suggesting that they were carried inside the EVs and being protected by the EV membrane. Intriguingly, the addition of 1% Triton X-100, which dissolved the EV membrane, together with proteinase K and RNase A treatment significantly reduced the level of let7b, but not that of Mir1 and miR34a (Fig. 3B), suggesting that these miRNAs possess differential resistance towards RNase A that was not attributed to the EV membrane.

In corroboration with this observation, our results showed that while RNase A treatment significantly reduced the amount of RNA extracted due to the removal of extracellular cell-free RNA, the presence of Triton X-100 did not further reduce the RNA quantity in testicular EVs (Fig. S6D), suggesting that the miRNA carried by testicular EVs is resistant to RNase A treatment. Of note, the RNase resistance requires the native RNA structure as alkaline hydrolysis or RNase A treatment after RNA extraction sensitized the candidate miRNAs to RNase A (Fig. S6E-F), suggesting that the miRNAs in testicular EVs possess specific resistance towards RNases in their native structures. These miRNAs were also carried by spermatozoa (Fig. 3C), suggesting the contribution of testicular EVs to the sperm RNA payload.

### Involvement of testicular EVs in spermatogenesis

Finally, we sought to investigate the involvement of testicular EVs in the development of germ cells. To achieve this, we injected GE4689, a small molecule that inhibits the release of EVs ^35^, into either the interstitial space or the seminiferous tubules via efferent duct injection. We performed the experiment with two-time points: a single injection followed by a 48-hours incubation (T1D2); and three injections (once every two days) and a total of 7-days incubation (T3D7). T3D7 injection was not feasible for the seminiferous tubules injection. Our results showed that a single injection with 48 hours incubation has no effect on the morphology and germ cell composition as compared to the vehicle control, regardless of the injection sites (Fig. 4A-B). Of note, a repetitive injection in the interstitial space and a longer incubation of 7 days led to a significant increase in the number of seminiferous tubules that showed signs of apoptosis. Intriguingly, apoptosis of Leydig cells and other interstitial cells located at the injection compartment were not observed (Fig. 4A-B). These results suggest that the testicular EVs, at least those in the interstitial space, were involved in normal germ cell development, likely via an inter-compartment communication.

**Fig. 4.**
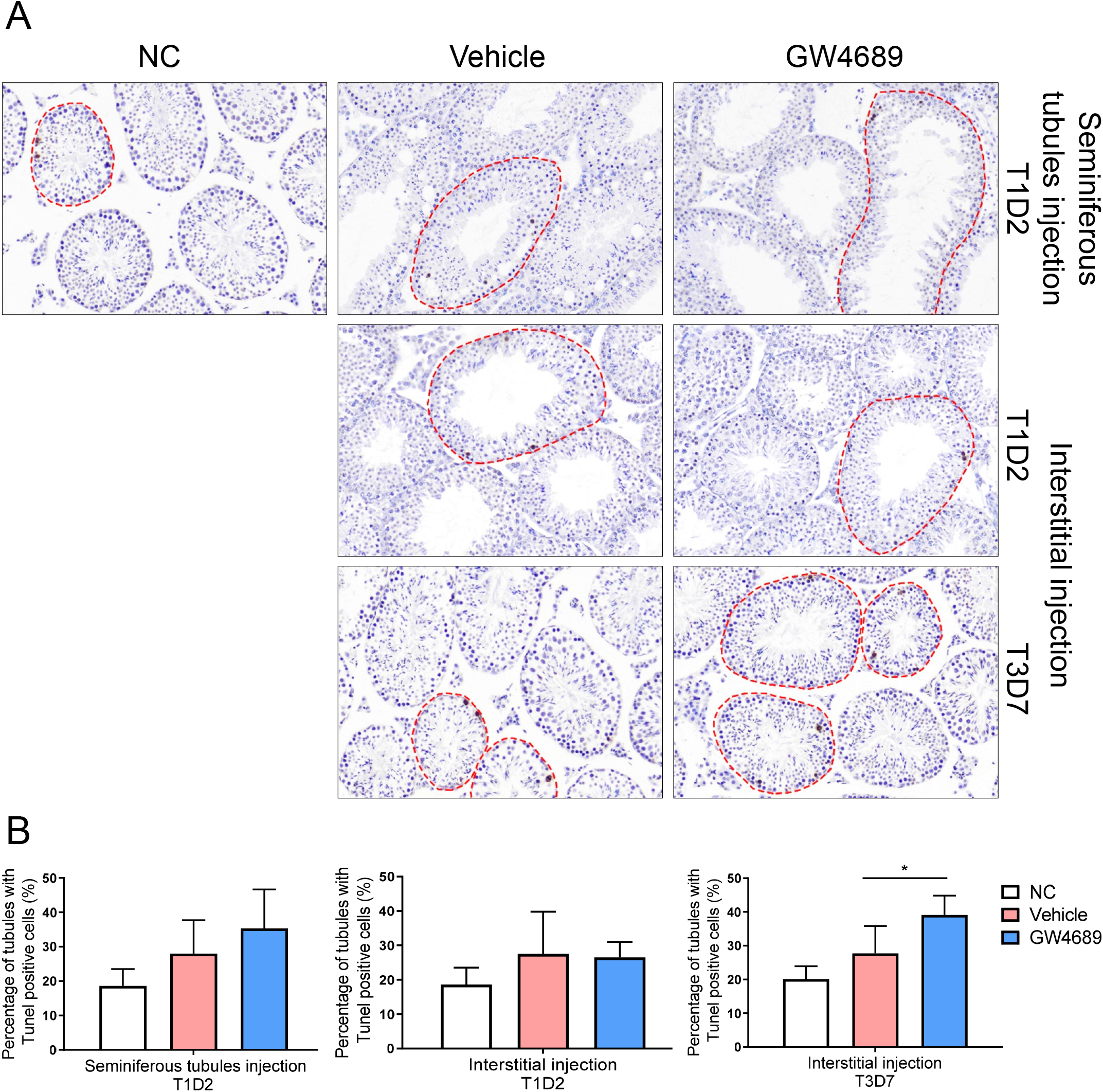
Inhibition of testicular extracellular vesicles elevated testicular germ cell apoptosis in mice. (A) Representative images of TUNEL assay performed on testes sections treated with GW4869 in the seminiferous tubules compartment (Top panel) or interstitium compartment (Middle and bottom panel) for the indicated days. T: treatment times; D: treatment days. Dashed line indicates the tubules with TUNEL positive cells. (B) Quantification from these images is shown. Data is presented as mean ± S.D. One-way ANOVA, Bonferroni post hoc test: * p <0.05.

## DISCUSSION

In this study, we have established a one-step tissue dissociation protocol that facilitated the isolation of testicular EVs with limited organelles contamination. Our protocol has omitted the use of trypsin which may disrupt the haploid germ cells/spermatozoa, leading to the release of intracellular membranous content that interferes with the characterization of EVs. Therefore, the proteomic and RNA-sequencing data presented here may represent a comprehensive profile of proteins and small RNAs with a low background interference. Our tissue dissociation protocol also preserves cell surface antigen which allows the isolation of a specific cell type for primary culture or for antibody-based characterization.

The present study is the first report on the characterization of testicular EVs. We showed that the testis microenvironment consists of microvesicles and exosomes which can be uptaken by the somatic cells and germ cells in both the interstitial space and seminiferous tubules, suggesting that the communication between these cells may involve EVs. Interestingly, while our proteomic data reveals two clusters of genes enriched in Leydig cells and macrophages, the cells in the interstitial space that uptake the testicular EVs. Since testicular EVs were capable of passing through the basement membrane and blood-testis barrier, we postulate that the EVs secreted by the Leydig cells and macrophages may act locally within the testis and modulate spermatogenesis in the seminiferous tubules. This is further supported by the effects of GW4869 in the apoptosis within the seminiferous tubules after being injected into the interstitial space. Unfortunately, a repetitive injection in the seminiferous tubules was not feasible and therefore the effects of testicular EVs without the seminiferous tubules cannot be investigated in the current study.

Besides, our result showed the presence of a non-negligible amount of microbial EVs and their RNA cargoes in the testicular microenvironment. While the function of the bacterial rRNA and tmRNA in testicular microbial EVs remain elusive, accumulating evidence has suggested a microbes-host communication where the small RNA in microbial EVs are internalized by the host cells and regulate cytokine production, signalling pathways and miRNA-like translational regulation ^36,37^. In fact, microbial RNA is present in sperm RNA and has been postulated to reflect the diversity of the associated bacteriome ^38^. These previous results and our findings collectively support the hypothesis that the sperm can internalize microbial EVs and carry the small RNA cargoes within. The involvement of these microbial EVs small RNAs in the regulation of spermatogenesis and in the epigenetic inheritance of microbiome warrant future investigation.

Our proteomic study revealed a plethora of proteins in the testicular EVs that have been implicated in male fertility, suggesting that the communication mediated by testicular EVs is required for spermatogenesis. Indeed, mouse genome informatics databases show that chemically induced mutation in the Smpd3 gene, which encodes for an enzyme required for EV release, reduced fertility. Also, gene knockout of Smpd3 shows delayed testis development and perturbs spermatogenesis at secondary spermatocytes. These results support an important role of testicular EVs in spermatogenesis. Surprisingly, although the Sertoli cell line and spermatogonial cell lines are capable of uptaking the EV, these cells did not release a measurable quantity of EVs in culture. It is known that the efficiency of in vitro spermatogenesis is very low. It is plausible that the absence of EV-mediated cell communication underlies the poor differentiation efficiency of germ cells in vitro.

Our results showed that testicular cells, including spermatogonia, Sertoli cells and sperm are highly efficient in uptaking testicular EVs but not EVs released by other somatic cells. This suggests the involvement of a specific targeting mechanism rather than a direct fusion of the membrane ^20^. Notably, around 10% of the proteins identified in the testicular EVs are involved in identical protein binding, suggesting a potential receptor-mediated uptake. Interestingly, our results also showed that sperm are highly efficient in uptaking the testicular EVs which may be a route to convey RNA into the transcriptionally inert sperm. Of note, the efficiency of sperm in uptaking testicular EVs (~84%) has surpassed that observed in epididymosome (~40%) after similar incubation time ^39^, suggesting that the testicular EVs may be a better tool in conveying cargoes to sperm.

EV is known to carry RNA cargoes that are protected from RNase by the lipid membrane. As expected, the candidate miRNAs in the testicular EVs were protected from RNase. Surprisingly, a portion of the miRNAs demonstrated another level of RNase resistance that requires a native conformation independent of protein-binding. The physiological significance of such RNase resistance warrants further investigation. Since the testicular EVs can deliver the cargoes to the head region of spermatozoa, we postulate that the stable miRNAs may be inheritable via the sperm epigenome and mediates paternal inheritance. Further mechanistic study on the RNase resistance of miRNAs may shed light on the synthesis of stable RNA molecules for gene therapy and genetic engineering.

Taken together, our study has successfully isolated the testicular EVs and characterized the protein and small RNA cargoes, providing valuable resources for future studies. We have further demonstrated its potential involvement in testicular cell communication and revealed a unique protection mechanism of the small RNA cargoes from RNases degradation, which may be involved in spermatogenesis and paternal inheritance via sperm.

## MATERIAL AND METHODS

### Animals

All animal experiments were approved by the Animal Research Ethics Committee of the Chinese University of Hong Kong (20-081-ECS). Eight-week-old male C57BL/6 mice were purchased from the laboratory animal service center (LASEC) of the Chinese University of Hong Kong.

### Testis dissociation

The mice were sacrificed by cervical dislocation. For the one-step method, seminiferous tubules were suspended in Accumax Cell Aggregate Dissociation Medium (20mg/ml) (Invitrogen; 00-4666-56) and incubated on a rocking platform at room temperature for 1 hour. The dispersed cells were filtered through a 40um cell strainer (Corning; 352340) to a 50ml falcon tube and washed with PBS. Filtered cells were pelleted at 300g for 5 minutes and the EV-containing supernatant was further filtered through using a 0.8um syringe filter (Millipore; SLAA033SB). The two-step double-enzymes digest method was performed as described ^23^, seminiferous tubules were suspended in PBS containing collagenase (1mg/ml) (Sigma-Aldrich) and incubated at 37°C for 10 minutes. The cells were washed once with PBS and further incubated in Trypsin (0.25%) (Gibco) and Dnase I (5mg/ml) (Sigma-Aldrich) at 37°C for 10 minutes. 10% FBS were added to inactivate the trypsin upon single-cell suspension and the dispersed cells were washed with PBS and Dnase I before filtered through using a 40um cell strainer. Filtered cells were pelleted at 300g for 5 minutes and all EV-containing supernatants were collected and filtered through using a 0.8um syringe filter.

### Isolation of EVs

For the one-step method, EV was isolated using exoEasy Maxi Kit (Qiagen, 76064) according to the manufacturer's instructions and further concentrated by Ultracentrifugation (Hitachi CS-150GXII Micro UITRAcentrifuge), performed at 100,000g for 90 minutes at 4°C.

For the two-step double-enzymes digest method, EV was isolated by differential centrifugation, performed first at 10,000g for 30 minutes at 4°C using Beckman Avanti J-E Centrifuge, then at 100,000g for 90minutes at 4°C using Beckman Coulter Optima XPN-100 ULTRAcentrifuge.

### Transmission electron microscope

Morphology of testicular EVs were observed using a transmission electron microscope. EVs from two testes were resuspended in 20 μl PBS and fixed with 2% paraformaldehyde until use. Ten μl of EVs were added onto the formvar grid (200 mesh) for 30-60 min and excess fluid was removed with filter paper. EVs were fixed with 1% glutaraldehyde for 10 min, followed by negative staining with 2% uranyl acetate for 2 min and images were captured using a Hitachi H-7700 transmission electron microscope.

### Dynamic Light scattering

Concentrated EVs were resuspended in 100 μl PBS and 50 μl of EVs were diluted in 950 μl PBS for determination of size distribution using the dynamic light scattering (Zetasizer Nano-ZS system). Three independent measurements were performed for each sample.

### Western blot

Protein lysate from EVs or cells was obtained by incubating with RIPA buffer with protease inhibitors. A total of 40 μg protein were electrophoresed under denaturing conditions on 12% polyacrylamide gels and transferred onto PVDF membranes. After blocking with 5% non-fat milk for 1 hour, membranes were immunoblotted with primary antibodies to CD 81, CD 63, Calnexin, Golgin 97, GAPDH overnight at 4 °C, followed by the relevant HRP-conjugated secondary antibodies incubation for 1 hour at room temperature. Bands were visualized by Prime ECL.

### Uptake of EVs

Concentrated EVs were labelled with PHK67 Green Fluorescent Cell Linker Mini Kit (Sigma-Aldrich, MINI67-1KT) according to the manufacturer’s instructions. EV concentrations were determined by protein assay. To study the EV uptake in testicular cells or somatic cells in vitro, indicated amount of labelled EVs or vehicle controls were incubated with C18-4, TM4, GC1-spg and 293FT cells when cells reached 60-70% of confluency in 24-well plates for different time-points or incubated with 300,000 sperms collected from mouse cauda epididymis and vas deferens in 60 mm dishes with 1 ml HTF medium for 4 hours. The uptake of EVs in cells or sperms were analyzed using immunofluorescence staining or flow cytometry. To study the EV uptake in mouse testis, labelled EVs or vehicle controls were injected to the seminiferous tubules or interstitium of mouse testes. After 24 hours, mice were sacrificed and the testes were collected. Snap frozen testicular sections (5 μm) were fixed with 4% paraformaldehyde for 15 min and slides were mounted in an antifade mounting medium with DAPI.

### Cell line

C18-4 cells were maintained in DMEM medium supplemented with 10% FBS, 1mM sodium pyruvate, 1% L-glutamine, 1 x nonessential amino acids at 35 °C with 5% CO2. GC1-spg cells and TM4 cells were cultured in DMEM or DMEM/F12 medium respectively containing 10% FBS at 37 ℃ with 5% CO2. C18-4, GC1-spg, TM4 cells were seeded on five 150 mm dishes containing 20 ml of cell medium with exosome-depleted FBS (Thermo Fisher, A2720803) and cell medium was collected for EVs isolation when cells were grown to 95% confluency.

### Thy1+ isolation with MACS

1×10^7^ cells from one-step and two-step double-enzyme digest methods were resuspended in 200ul 2% FBS DMEM and incubated with 20ul primary antibody (1:10, Biotinylated Thy1.2 CD90.2) on ice for 15 minutes on a slow-rocking platform. MS columns (Miltenyi Biotec, 130-042-201) were first calibrated with 500ul 2% FBS DMEM. Cells were washed with PBS and resuspended with 500ul 2% FBS DMEM before loaded into the column to allow Thy1-cells to flow through by gravity. Columns were washed twice with 500ul 2% FBS DMEM and finally, Thy1+ cells were eluted with 1ml 2% FBS DMEM.

### Flow cytometry

Cells collected from one-step and two-step double-enzyme digest methods were washed once with PBS and permeabilized overnight with 70% ethanol at −20°C. Samples were stained with 40ug/ml PI in PBS and 100ug/ml Rnase A and incubated at 37°C for 30 minutes in dark. PI stained cells were analyzed with BD LSR Cell Analyzer, counting 10,000-20,000 events per sample.

Flow cytometry was used to determine the percentage of EV uptake in cells or sperms. After the incubation period with labelled EVs or vehicle control, cells were washed with PBS once and trypsinized, pelleted by centrifugation at 1,000 rpm for 5 min and resuspended in 300 μl of cell medium, while sperms were washed with PBS once, pelleted by centrifugation at 1600 rpm for 5 min and resuspended 500 μl in PBS. FITC positive cells or sperms were analyzed BD LSR Fortessa Cell Analyzer, counting 50,000 cells or sperms for each sample.

### Immunofluorescence staining

Cells were fixed in 4% paraformaldehyde for 15 min, permeabilized with 0.1% Triton X-100 for 10 min followed by incubation with Alexa Fluor 568 Phalloidin (dilution 1:200) for 20 min. Cells were counterstained with Hoechst 33342 (1:2000) for 10 min.

### Alkali/detergent and nuclease treatment of testis EV

Prior to RNA extraction, EV/sperm samples were treated with or without 1% Triton X-100 (Boehringer Mannheim, 1332481), incubated at room temperature for 30 minutes to disrupt the membrane. The samples were then treated with Proteinase K (0.05ug/ul) (Qiagen, 1014023) at 37°C for 10 minutes, inactivated with PMSF (5mM) (Sigma-Aldrich, P7627) at room temperature for 10 minutes then treated with Rnase A (0.05ug/ul) (Thermo Scientific™, EN0531) at 37°C for 20 minutes. Alkaline hydrolysis was performed as described in (Lemire et al., 2016). Briefly, samples were first incubated at 65°C for 1 hour in 0.1M Tris (pH 8.0) and 1M NaOH, then neutralized with 1M HCl at a ratio of 2.33:1:1.

### RNA extraction, reverse transcription and real-time PCR

Total RNA was extracted using the miRNeasy Mini Kit (Qiagen, 217004) according to the manufacturer's instructions. 10pg of mirVana miRNA mimic miR-199a-5p (ThermoFisher, Assay ID MC10893 and 002304) was added to 0.1-0.5ug of total RNA as a spike-in control for RT-qPCR normalization. Reverse transcription was performed using PrimeScript™ RT Master Mix (TaKaRa, RR036A) for novel miRNA and TaqMan™ MicroRNA Reverse Transcription Kit (Applied Biosystems, 4366597) for the candidate miRNAs let-7b-5p (ThermoFisher, Assay ID 000378) and miR-34a-5p (ThermoFisher, Assay ID 000426). 1ul of cDNA was used for real-time PCR using TaqMan™ Universal PCR Master Mix (Applied Biosystems, 4364340) on the ABI QuantStudio 7 Flex Real-Time RCR system. Primers and probes used are listed in the Supplemental Table S3. Results were calculated with 2(−ΔΔCt).

### Small RNA sequencing

EVs RNA after treatment with RNase A and Proteinase K was extracted as described above and used for small RNA-Seq which was performed by BGI (Shenzhen, China). Small-RNA libraries were prepared, and the PCR products were sequenced using BGISEQ-500 technology to obtain the indicated number of reads. After elimination of low-quality reads, clean reads were mapped to sRNA database such as miRbase, Rfam, siRNA, piRNA, snoRNA and etc. The proportion of all kinds of annotated sRNAs were analyzed. Novel sRNAs were predicted from those unknown tags.

### LC-MS/MS

Testicular EVs protein were extracted with RIPA buffer and LC-MS/MS proteomics analyses were performed by Winninnovate Bio (Shenzhen, China). Gene Ontology (GO) enrichment and KEGG pathway enrichment were analyzed.

### GW4869 Treatment

To study the effect of exosome generation inhibitor GW4869 (Sigma-Aldrich, D1692) on spermatogenesis in mouse, GW4869 dissolved in DMSO (20 μM, 20μl/testis) or vehicle were injected into the mice seminiferous tubules or interstitium of testes once or every other day for three times under full anesthesia with 100mg/kg ketamine and 10 mg/kg xylazine mix. After the indicated treatment time, mice were sacrificed and testes were fixed in 4% paraformaldehyde, dehydrated, and embedded in paraffin. The TUNEL assay was performed on the testis sections (5 μm) with ApopTag plus peroxidase in situ apoptosis detection kit (Millipore, S7101).

### Statistical analysis

Statistical analysis was carried out using GraphPad Prism version 8.02. One sample t-test was used to compare group means. Ordinary One-way Analysis Of Variance (ANOVA) with Dunnett's post hoc test was used for analysis involving three or more groups of samples. A P value of <0.05 was considered significant.

## ACKNOWLEDGEMENT

We would like to thank Prof Simon Wing for providing the C18-4 cell line.

## AUTHOR CONTRIBUTIONS

KHKC, SYC, WL and EKLF conceived and designed the experiments; KHC, WL, SYC, JJ, ZT performed the experiments and analyzed the data; SS, WW, HC, GX, HCY and LL gave intellectual advice; EKLF provided funding support; KHKC, SYC and EKLF wrote the paper.

## DATA AVAILABILITY

Raw data generated in this study have been deposited in the database. Proteomic data is available on the PRIDE database (Project accession: PXD021766). Small RNA sequencing data is available on the GEO database (Accession number: GSE158402).

## Supplemental data file

**Supplemental Table S1| List of proteins identified in testicular EVs.** Table listing the proteins commonly found in two individual batches of testicular EV samples (n>3 mice in each batch). Confidence coefficient (−10lgP) and number of unique peptide mapped (#unique) are provided for each entry. Samples are ranked by the average number of unique peptides.

**Supplemental Table S2| List of small RNAs identified in testicular EVs.** Table listing the small RNA commonly found in two individual batches of testicular EV samples. The counts and transcripts per million (TPM) are shown for each entry. Samples are ranked by the average number of counts.

**Supplemental Table S3| Oligo information.**

## Supplemental figure legends

**Supplemental Fig. S1.**
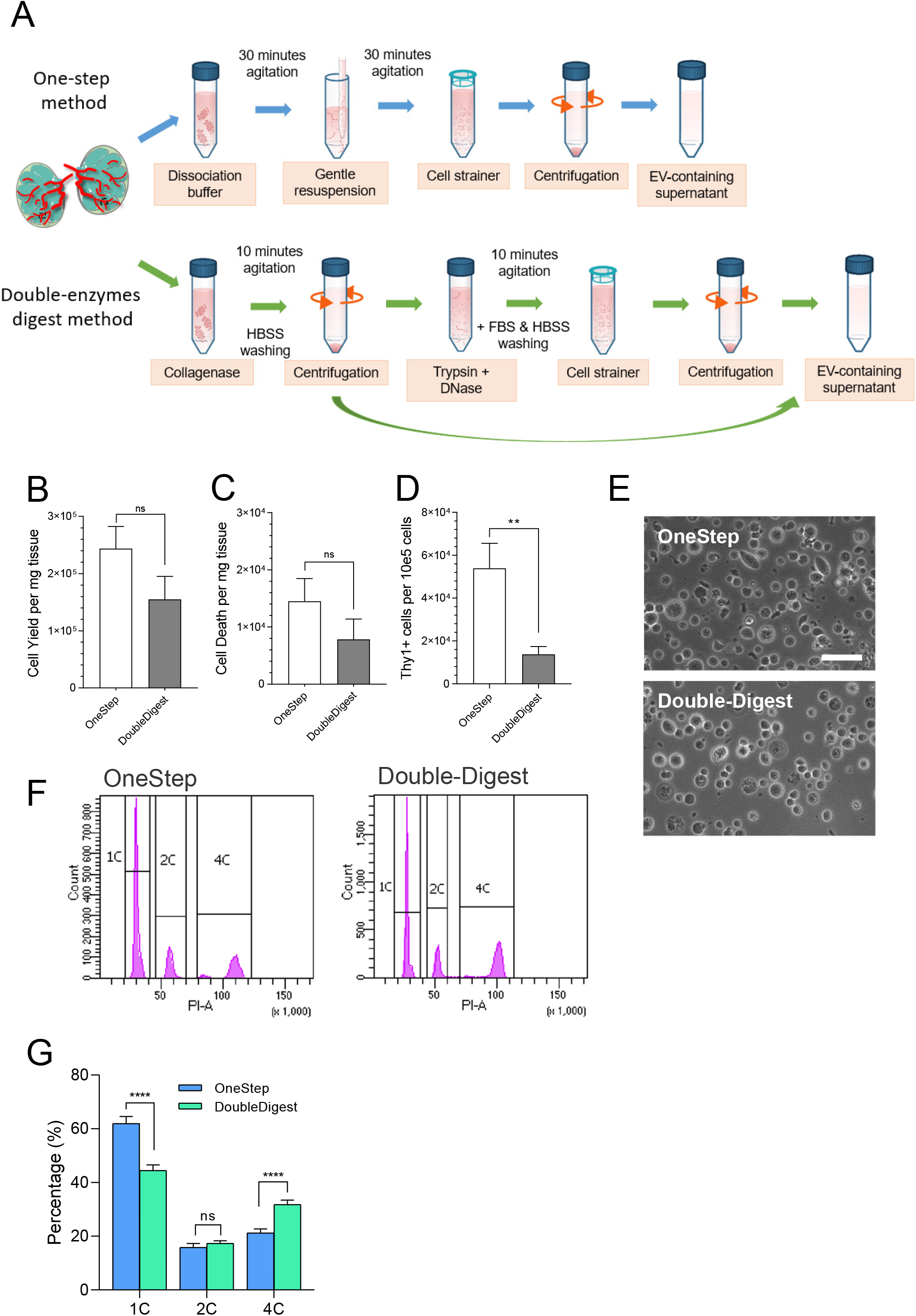
Comparison of one-step and double-enzyme digestion methods for mouse testis dissociation. (A) Schematic diagram comparing the one-step testis dissociation method and the two-step double-enzyme digestion method. (B-C) Total cell yield (B) and cell death (C) per mg tissue extracted from indicated methods. (D) Cell count of Thy1+ cells isolated by magnetic activated cell sorting (MACS) from testicular cell suspension obtained from both methods. (E) Representative phase contrast imaging of single cell suspension obtained by indicated methods. Scale bars = 50um. (F) Flow cytometry analysis showing the composition of PI-stained cells obtained from indicated methods. (G) Quantification of DNA ploidy (haploid - 1C; diploid - 2C; tetraploid - 4C) by flow cytometry in testicular cell suspension prepared by both methods.

**Supplemental Fig. S2.**
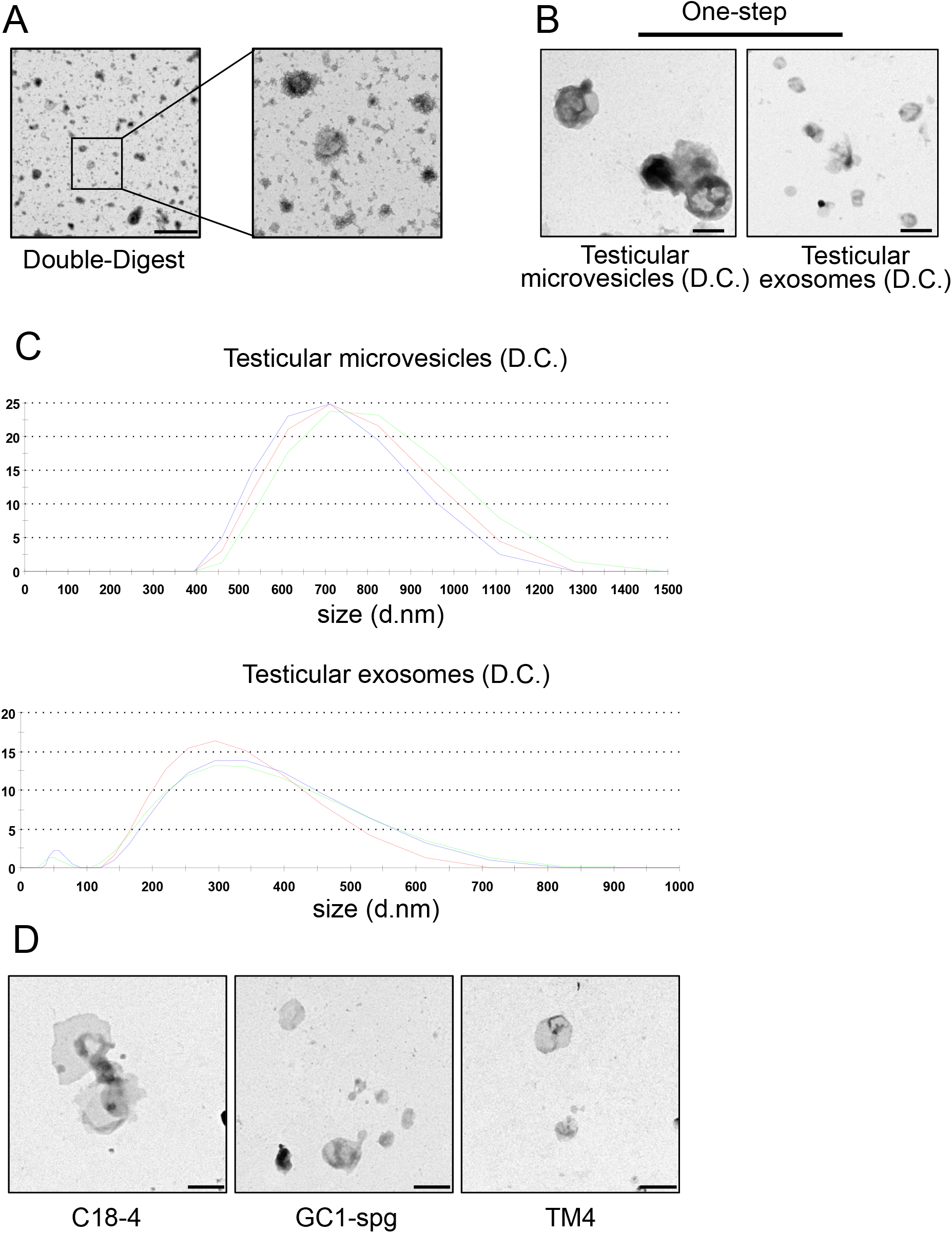
Isolation and characterization of extracellular vesicles in mouse testis and testicular cell lines. (A) Representative transmission electron microscopy images of testicular EVs isolated from testicular cell suspension prepared by double-enzymes digestion method followed by affinity column isolation. Note the presence of debris in the background and surrounding the cup-shape EVs. Scale bar: 500 nm. (B) Representative transmission electron microscopy images of testicular microvesicles and exosomes isolated by differential ultracentrifugation (D.C.). Scale bar: 500 nm. (C) Size distribution of testicular microvesicles (upper panel) and exosomes (bottom panel) isolated from one-step testis dissociation followed by differential centrifugation as determined by dynamic light scattering. (D) Representative transmission electron microscopy images of EVs isolated from the conditioned medium of spermatogonia cell lines (C18-4 and GC1-spg) and Sertoli cell line (TM4) by differential centrifugation. Scale bar: 500 nm.

**Supplemental Fig. S3.**
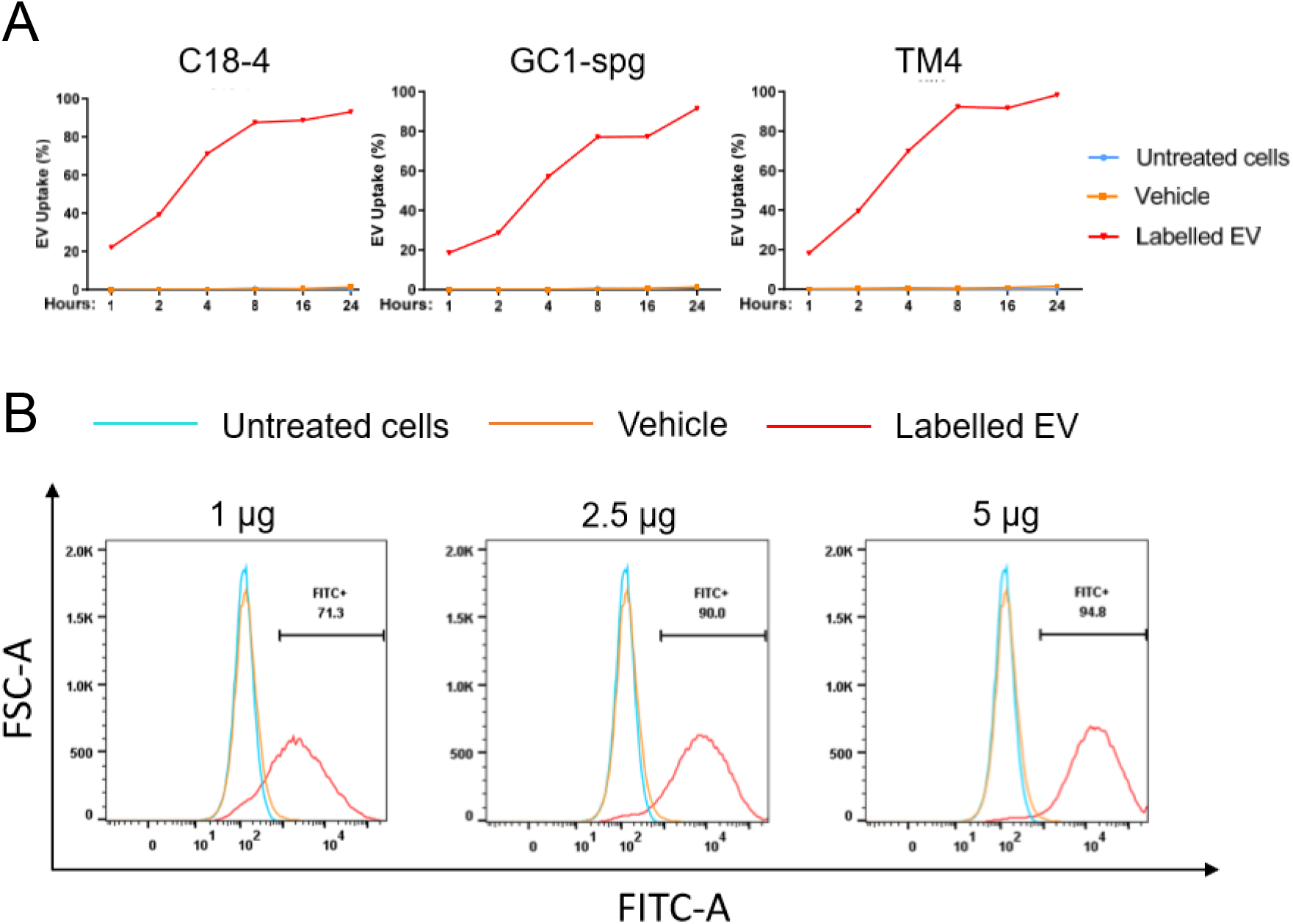
Time and dose-dependent uptake of testicular extracellular vesicles by testicular cell lines. (A) Quantification of the percentage of indicated cell lines that uptake the PKH-labelled testicular EVs at indicated time points as examined by flow cytometry. (B) Flow cytometry analysis of C18-4 cell after treatment of indicated amount of testicular EVs (in terms of protein amount) for 24 hours.Diagram showing the percentage of microbial rRNA mapped to the indicated microbes.

**Supplemental Fig. S4.**
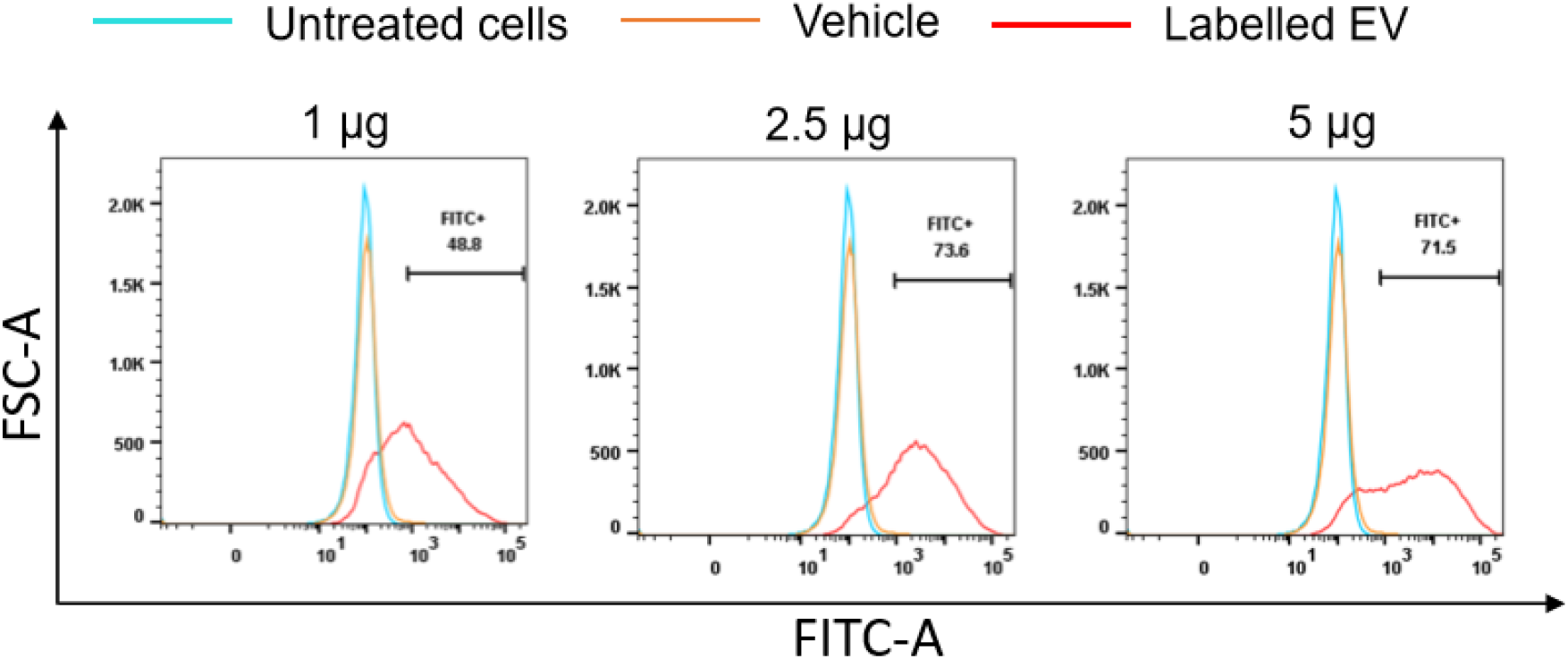
Uptake of extracellular vesicles secreted from HEK293 cells by C18-4 cells. Flow cytometry analysis of C18-4 cell after treatment of indicated amount of EVs isolated from condition medium of HEK293 cells (in terms of protein amount) for 24 hours. Note the plateau in the percentage of FITC+ cells in 2.5 and 5 ug EV treatments.

**Supplemental Fig. S5.**
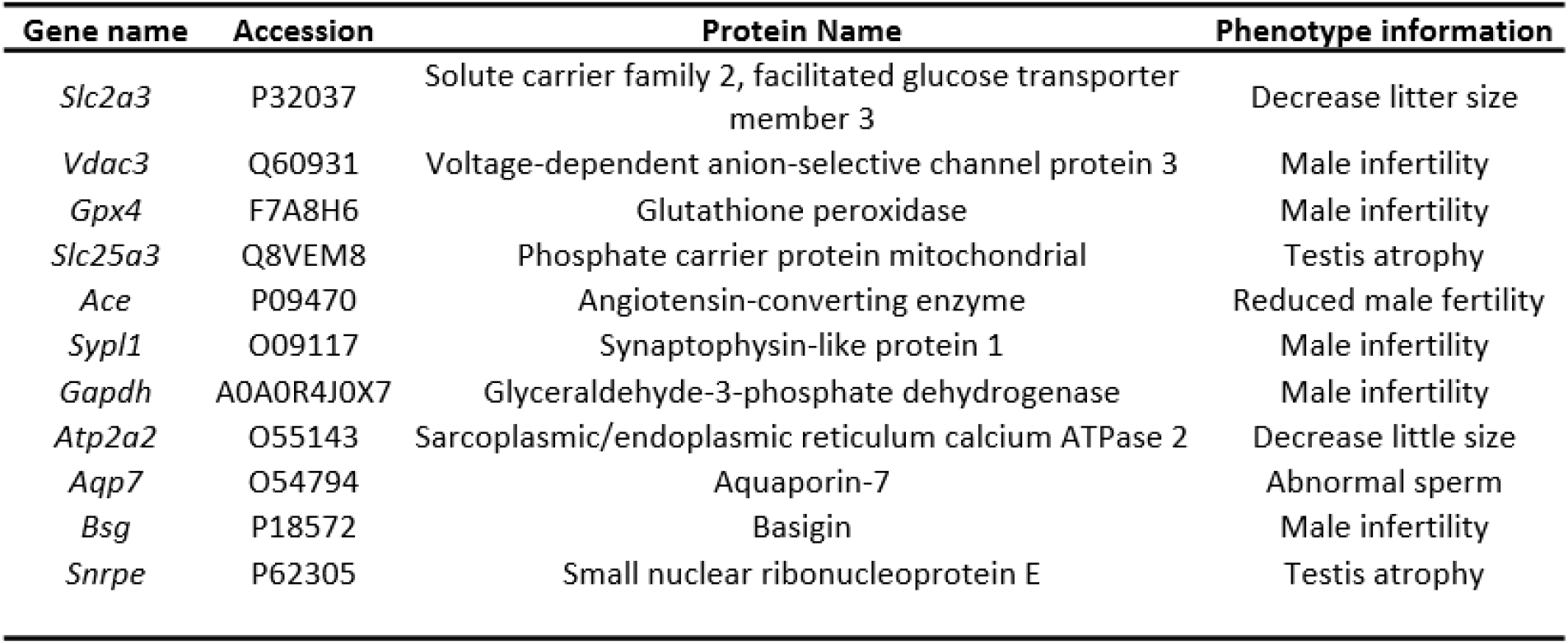
Phenotype of male mice with knockout of genes encoding top proteins identified in testicular EVs. Phenotype information of genes encoding the proteins with the highest number of unique tags in the testicular EVs. Phenotype information is obtained from Mouse Genome Informatics (MGI) database.

**Supplemental Fig. S6.**
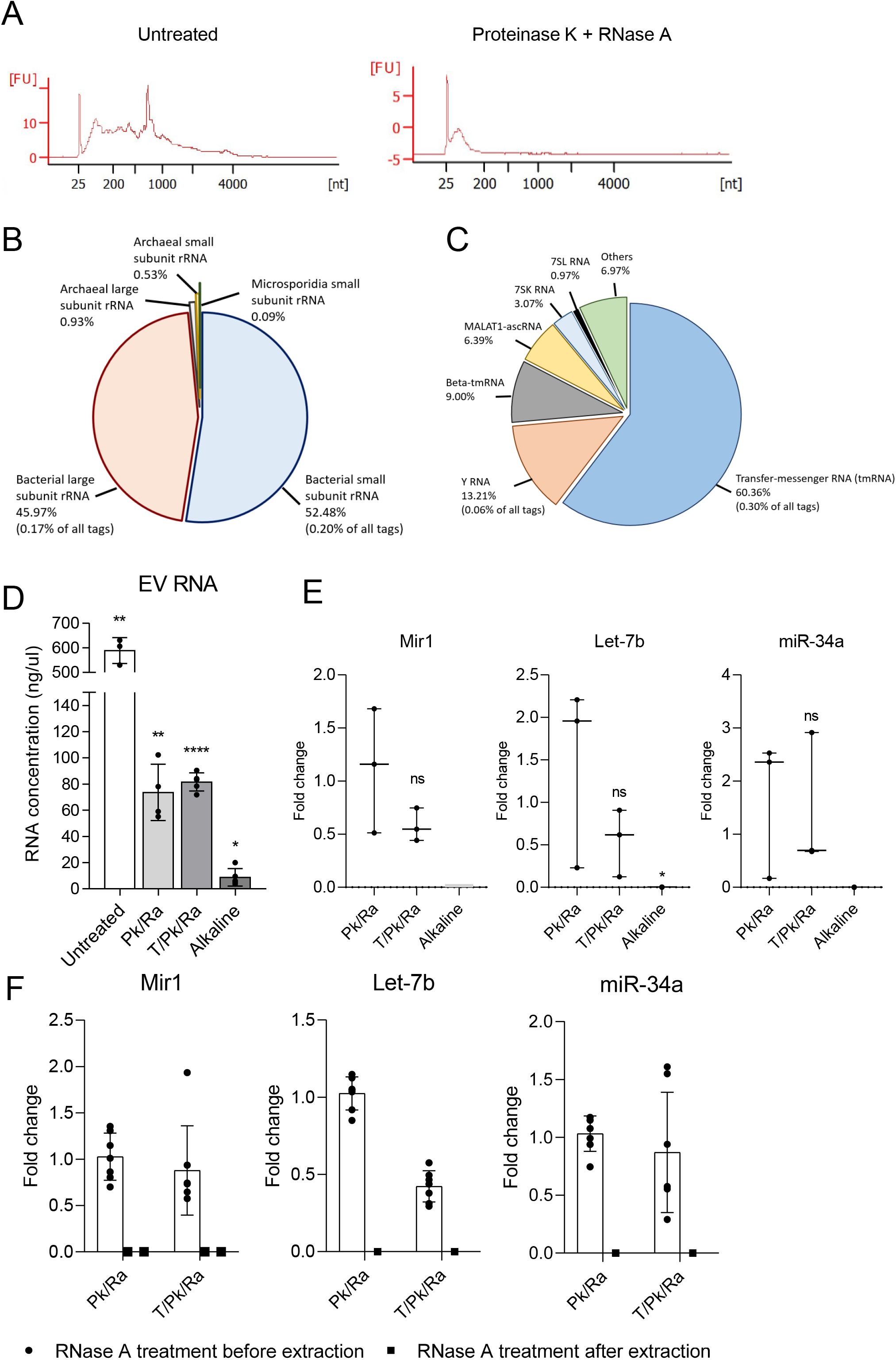
Characterization of the small RNA cargoes of testicular EVs. (A) Bioanalyzer traces showing the size distribution of RNA cargo carried by testicular EVs before (left panel) and after Proteinase K and RNase A treatment (right panel). (B) Diagram showing the percentage of microbial rRNA mapped to the indicated microbes. (C) Diagram showing the percentage of top 6 most abundant sncRNA in testicular EVs based on Rfam database. (D) Quantity of RNA extracted from testicular EVs without treatment (untreated), with proteinase K and RNase A treatment (Pk/Ra), with Pk/Ra treatment in the presence of 1% TritonX-100, or with alkaline hydrolysis treatment. (E) Realtime PCR results showing the level of novel Mir1, let-7b and miR-34a in testicular EV samples with proteinase K and RNase A treatment (Pk/Ra) in the presence or absence of 1% TritonX-100 (T) or with alkaline hydrolysis. Same volume of extracted RNA was used for reverse transcription due to the extreme low quantity of RNA in the alkaline hydrolysis group. (F) Realtime PCR analysis of Mir1, Let-7b and miR-34a with additional RNase A treatment after RNA extraction. Data is presented as mean ± S.D. ** p <0.01, **** p<0.0001, ns - not significant.

